# Cytochrome *c* oxidase I deep amplicon sequencing for metabarcoding of equine strongyle communities: unexpectedly high *Strongylus* spp. burden in treated horses

**DOI:** 10.1101/2025.05.12.653446

**Authors:** Jürgen Krücken, Irina Diekmann, Sandro Andreotti, Christina M. Bredtmann, Susan Mbedi, Sarah Sparmann, Jennifer S. Schmidt, Fernando de Almeida Borges, Mariana Green de Freitas, Guillaume Sallé, Heribert Hofer, Jacqueline B Matthews, Thomas Tzelos, Martin K. Nielsen, Tetiana A. Kuzmina, Georg von Samson Himmelstjerna

## Abstract

Equines are parasitized by complex communities of Strongylidae (Nematoda) comprising multi-species infections. Currently, the Cyathostominae are most prevalent, while the *Strongylus* species are only rarely detected. Since eggs and, in most cases, infective larvae cannot be differentiated to species level, with the exception of *Strongylus* spp., species-specific knowledge of the pathology, epidemiology and ecology of these parasitic nematodes is limited. Reference sequence data for several cyathostomin species are limited or missing. Deep amplicon sequencing of internal transcribed spacer 2 (ITS-2) regions of nematodes has been used in equines previously, although barcoding studies demonstrate a better species resolution for the cytochrome *c* oxidase subunit I (COI) region. The present study introduces a nemabiome method based on the sequencing of COI fragments. This method was applied to compare third stage larvae, representing strongyle communities, derived from regularly treated (RT) and never treated (NT) equine populations from Brazil, France (only RT), Germany, Ukraine, the UK, and the USA. Samples were predominantly from horses, but some were obtained from Przewalski’s horses (Ukraine), donkeys (Germany, Ukraine) and kulans (Ukraine). Most sequence reads (87.7%) were identified to the species level, but unclassified reads occurred more frequently in donkeys and kulans than horses. No obvious difference in species diversity and richness was observed between RT and NT equines. However, there were significant differences in species composition between the RT and NT groups. While *Strongylus* spp. were significantly more abundant in the NT groups, *Cylicocyclus nassatus*, *Cylicostephanus longibursatus,* and *Cyathostomum catinatum* were more abundant in the RT group, suggesting that strongyle communities in domestic equines may have been shaped by anthelmintic treatments in the last decades. The decreased classification success for reads from non-caballine equines suggests that there are more strongyle species specific for this rarely-investigated group and that additional efforts are needed to improve the sequence database, particularly for these hosts.

**Author summary:** This study shows that long-term deworming treatments have influenced strongyle nematode communities in equines. Our findings showed that regular deworming does not always reduce species richness and diversity. Noteworthy, we observed that the more pathogenic species such as *Strongylus vulgaris* and *Strongylus edentatus* were still present but in low abundance in equines. Due to their low abundance, less sensitive diagnostic methods, such as morphological examination of larval cultures might not detect these species, which would lead to an underestimated threat in equine herds. We applied an effective metabarcoding approach based on a reliable gene marker region to accurately detect these and other strongyle species in equines. The detection of various species was more effective for horse samples than for those from donkeys and kulans. An expansion of the current database that includes more specimens from more rare species obtained from different equine species can improve identification and understanding of these complex multi-species communities in the future. In summary, the study underscores the importance of continuing monitoring equine herds based on sensitive methods such as metabarcoding to evaluate the current nematode communities and to adapt and develop treatment strategies for managing strongyle infections in equines.

## Introduction

Although most grazing mammals are parasitized by multiple nematode species of the order Strongylida, the situation in equines is quite special. While representatives of the Molineoidea superfamily (for example, *Nematodirus* spp.) are not relevant to equids and only one species of the superfamily Trichostrongyloidea, *Trichostrongylus axei*, parasitizes equids, members of the superfamily Strongyloidea are predominant. The subfamily Strongylinae includes 14 equine species in five genera (1), including *Strongylus vulgaris* and other members of the *Strongylus* genus, which are important due to their high pathogenicity caused by the migratory behavior of their larval stages (2–4). Less pathogenic to an extent, but ubiquitously occurring in equines, are the Cyathostominae, also grouped as cyathostomins (5). A unique feature of the subfamily Cyathostominae is its high species diversity. The total number of recognized species of Cyathostominae is 50 (1), but several studies using molecular data suggest that the actual number of species might be higher due to the presence of morphologically undistinguished species that differ in their genotype, so-called cryptic species (6–10).

Species identification and taxonomy of cyathostomins are hampered by morphological similarities between some species and a lack of features that act as informative phylogenetic traits (11). Globally, only a few experts exist that can reliably identify adult cyathostomins to species level, and the morphological identification of eggs and for most species of infective third stage larvae (L3) is not possible. The fact that cyathostomin communities can be extremely diverse and mixed infections are the rule, complicates the epidemiological picture. In many horses, five to ten species are usually found, but the number can be as high as 25 cyathostomin species observed in a single horse if the number of worm specimens used for identification is sufficient (12). In a meta-analysis, Bellaw and Nielsen (13) identified three Cyathostominae species that were found at high prevalence and abundance in studies from seven regions located on five continents: *Cylicocyclus nassatus*, *Cylicostephanus longibursatus,* and *Cyathostomum catinatum*. Another five species were classified as moderately prevalent and abundant, while other species were assigned only with low or very low abundance. The diversity of cyathostomin communities, together with the limitations concerning species-specific identification, as well as the lack of any single species isolates, means that there is barely any information available regarding species-specific epidemiology, ecology, pathology, or spread of anthelmintic resistance (11).

Deep amplicon sequencing has been used to characterize species composition of various ecosystems using environmental DNA or samples enriched for certain types of organisms in a metabarcoding approach (14). Characterization of microbiota using deep amplicon sequencing based on partial or nearly full-length 16 S rRNA gene PCR products is the most widely used application of this type of approach. In eukaryotes, PCR’s targeting partial 18S rRNA and cytochrome *c* oxidase I (COI) genes, as well as partial and complete internal transcribed spacer (ITS) regions, have been used for metabarcoding. These markers differ in the speed of their evolution and in their ability to discriminate between taxa (15). Since prokaryotic 16S rRNA gene sequences are highly conserved, they are typically not suitable to identify specimens to species level, at least as long as short, partial sequences are used, as is necessary if Illumina sequencing technology is applied (16). In contrast, ITS-2 and COI have the potential to enable identification to species level using short amplicons (15). In ruminants trichostrongyle parasites such as *Cooperia* spp. and *Marshallagia* spp., ITS-2 is reported as an excellent marker to identify sequences at least to the genus level (17, 18). At the species level, different species of the same genus often differ in a very small number of base pair positions, as shown for *Haemonchus*, *Cooperia,* and *Marshallagia* species, and often the polymorphisms are not fixed between species. Thus, there is often no clear barcoding gap if ITS-2 sequences are used to discriminate between different members of the same genus (17, 18). For equines, the situation is more complicated due to there being (i) over 64 strongyle species potentially present, (ii) no sequence data available for roughly half of these species (19), (iii) the presence of cryptic species for several cyathostomins (6–10, 19), (iv) some species, for example, *Coronocyclus coronatus*, have been shown to possess two ITS-2 sequence versions that differ in size, and (v) morphologically distinct species, for example, *Cor. coronatus* and *Cylicostephanus calicatus*, can have identical ITS-2 sequences (6). Some of these features lead to misassignment of amplicon sequence variants (ASVs) to the wrong species.

In both ruminant and equine strongyle parasites, the barcoding gap is wider for COI sequences. They show high diversity at codon position three, often close to substitution saturation, which leads to only 90–95% sequence identity between closely related species (10, 19); for example, COI sequences are easily able to discriminate *Cor. coronatus* and *Cys. calicatus* (6). In comparison to ITS-2 sequences, COI sequences have the advantage that polymorphisms are more or less evenly distributed along the sequence length and do not occur in clusters. Therefore, reliable alignments can be easily obtained by aligning the translated protein sequences (18, 19).

Gastrointestinal nematode communities of large and small ruminants have been characterized in a nemabiome approach using deep amplicon sequencing of ITS-2 PCR products (20–24) by modifying a PCR to identify nematode species (25). The method has been used to identify species exhibiting anthelmintic resistance (21, 26–29). Most studies have been conducted using the Illumina MiSeq system; however, some have used PacBio sequencing technology (28, 30), arguing that the latter shows less bias regarding read frequency and amplicon length. The ITS-2 based approach has been used to characterize strongyle communities in horses from Australia (31, 32), Canada (33, 34), Sweden (30, 35), Thailand (36), the UK (37–39), and the USA (33, 40). Furthermore, this approach was utilized to assess strongyle community structure in zebras (41) and to study the effects of anthelmintics (42, 43) or treatment strategies (30). Courtot, Boisseau (44) compared ITS-2 and COI based metabarcoding approaches using various mock communities and sample types (eggs, larvae, adults); the COI barcode proved suboptimal in terms of higher PCR amplification bias, reduced sensitivity to detect rarer species and a higher divergence when comparing the expected community composition with the sequencing results. However, the COI PCR used did not overlap with the one used in previous projects established for meta-barcoding (6, 7, 10, 19). The latter were part of a project to generate ITS-2 and COI sequence databases from individual strongylid specimens morphologically identified by current experts. By obtaining molecular data for both target regions from the same worm and using multiple individuals per species, the derived databases were assessed for reliability and ability to detect cryptic species indicated by high diversity in COI sequences compared with identical clustering in ITS-2 sequences (6, 7, 10, 19).

Here, the suitability of a medium-long COI amplicon (653 ± 3 bp) to characterize strongyle community structures in samples from different countries and host species was explored using an amplicon previously assessed for taxonomic and phylogenetic purposes (6, 7, 10, 19). The size prevents sequencing of complete amplicons using available Illumina sequencing technologies, so, compared to traditional deep amplicon sequencing approaches that rely on considerable overlap of forward and reverse reads, modified bioinformatic pipelines needed to be implemented. This study aimed to assess incomplete sequencing of COI amplicons for providing reliable results for metabarcoding of equine strongyle communities, including the detection of closely related cryptic species. The method was applied to mock communities and then used to determine differences in strongyle community structures in horses from herds that have been regularly anthelmintic treated (RT) or those that are rarely/never treated (NT).

## Results

### Comparison of cytochrome *c* oxidase I PCR amplification efficacy between strongyle species

Efficacy of the first PCR was evaluated using plasmids containing inserts representing the amplicons from 12 strongyle species. After calculating efficacy for each of the three replicates from the exponential amplification plots, efficacies were plotted for each species (Fig 1). A One-way ANOVA did not provide any evidence that there might be differences in efficacy between species (p = 0.963). Visual inspection of the dataset revealed a similar spread of efficacy in individual replicates. The overall range for all specimens was 1.803 – 2.005, while the lowest efficacy for replicates in each species varied between 1.803 and 1.906, and the highest efficacy between 1.919 and 2.005. The mean efficacies ranged from 1.884 to 1.951.

**Fig 1.**
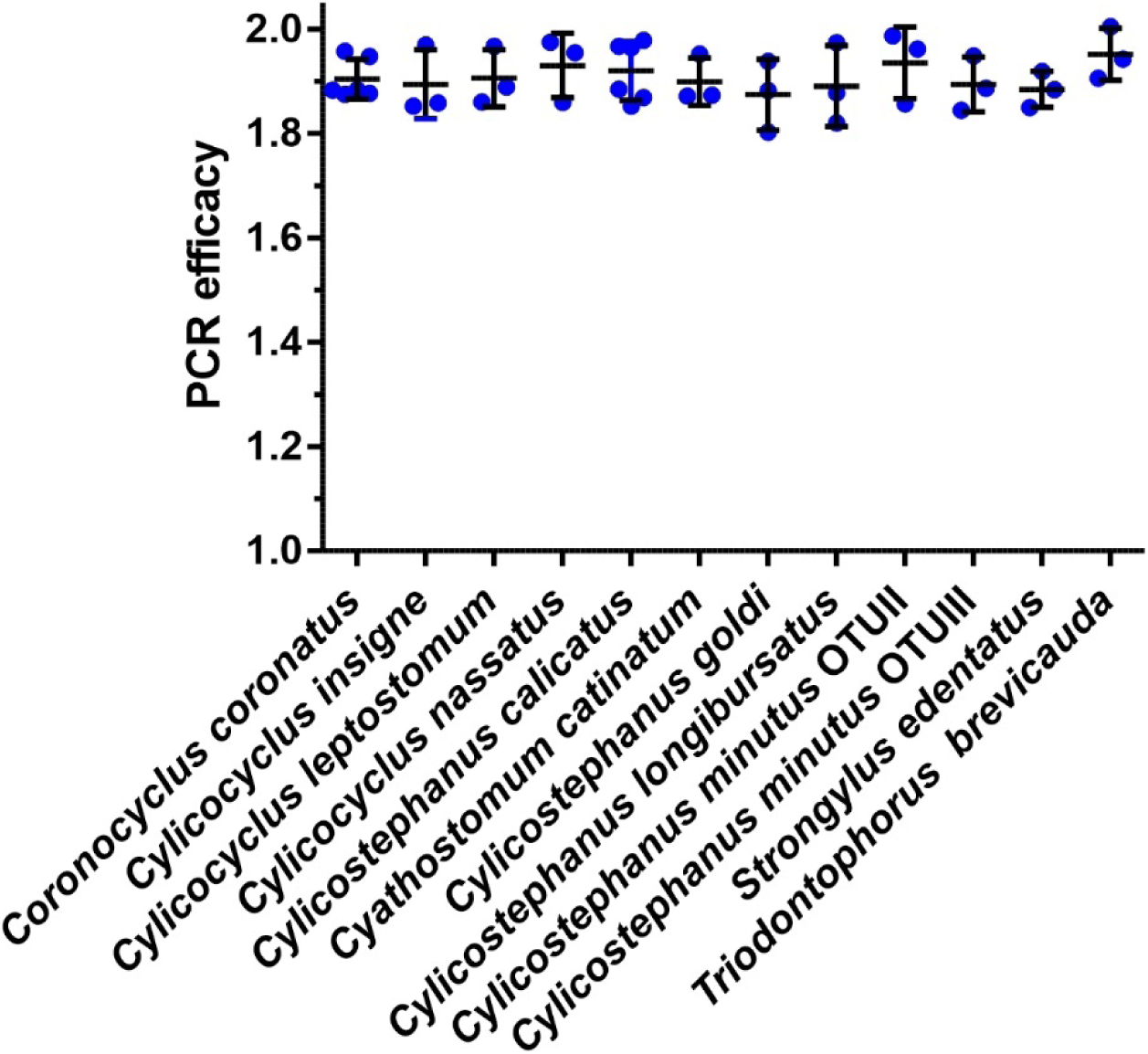
Cytochrome *c* oxidase I PCR efficacy of individual reactions for plasmids representing 12 strongyle species. For each species, three or six representative plasmid DNAs containing an insert with the PCR amplicon were used as templates for the PCR. Each replicate (blue dots) and means ± SD were plotted. One-way ANOVA revealed no evidence for differences between means (p = 0.878). Efficacies in individual PCR reactions were largely overlapping for all species.

### Sequencing of mock communities obtained by pooling of plasmid DNA

A mock community was created by equimolar mixing of plasmids encoding partial COI regions from 27 different strongyle species (*Coronocyclus* (*Cor.*) *coronatus*, *Cor. labiatus*, *Cor. labratus*, *Cor. sagittatus*, *Craterostomum* (*Cra.*) *acuticaudatum*, *Cyathostomum* (*Cya.*) *catinatum*, *Cya. pateratum*, *Cya. tetracanthum*, *Cylicocyclus* (*Cyc.*) *ashworthi*, *Cyc. elongatus*, *Cyc. insigne*, *Cyc. leptostomus*, *Cyc. nassatus*, *Cyc. radiatus*, *Cylicodontophorus* (*Cyd.*) *bicoronatus*, *Cylicostephanus* (*Cys.*) *bidentatus*, *Cys. calicatus*, *Cys. goldi*, *Cys. longibursatus*, *Cys. minutus* OTU II, *Gyalocephalus* (*Gya.*) *capitatus*, *Petrovinema* (*Pet.*) *poculatum*, *Poteriostomum* (*Pot.*) *imparidentatum*, *Strongylus* (*Str.*) *vulgaris*, *Triodontophorus* (*Tri.*) *brevicauda*, *Tri. serratus*, *Tri. tenuicollis*) to estimate the variation introduced by the deep amplicon sequencing method. The mock community sample was sequenced in parallel with the field samples obtained from equines. In total, 79,041 merged reads were obtained, of which none were removed by blast and abundance filtering, and the remaining available reads were included in the analysis. As shown in Fig 2, all 27 species were detected in the sample. The frequency at which individual species were found ranged from 0.27% - 10.3% of all reads (median 3.6%). The mean frequency of reads per species was 4.0% ± 3.1 (mean ± SD); this was not significantly different from the expected value of 3.7% (p = 0.642, one sample t test; p = 0.788, Wilcoxon signed rank test). Almost all species were represented by only a single ASV. The exception was *Cyc. leptostomus*, for which two ASVs were found, one with a frequency of 1.4% and one with a frequency of 0.001%, possibly caused by a PCR or sequencing error. There was also one ASV representing *Cys. minutus* OTUIII (0.01%), although no plasmid for this species was included in the mixture, and two ASVs that could not be classified to species level (both with frequencies of 0.01% and 0.01%).

**Fig 2.**
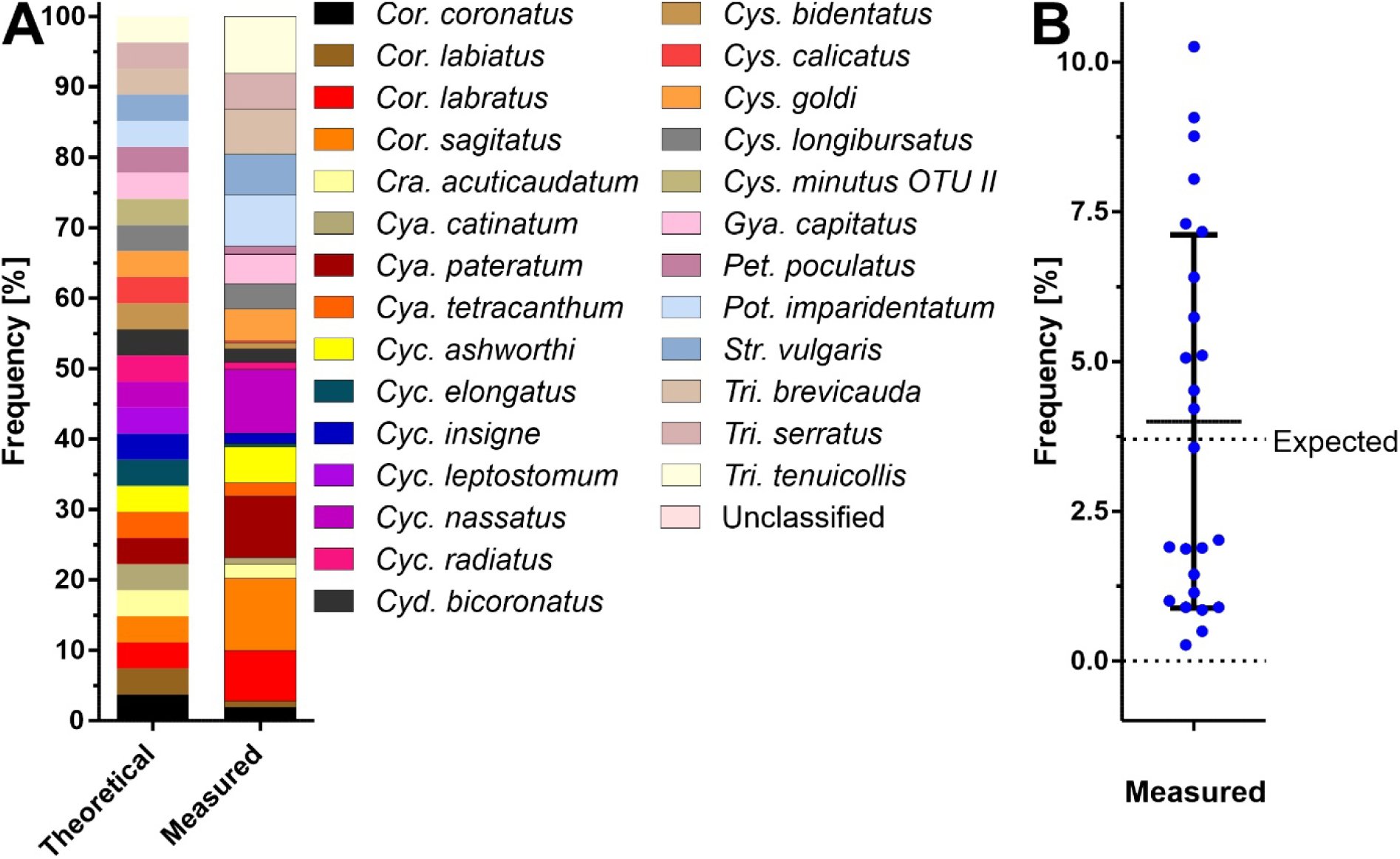
Comparison of the theoretical composition of a mock community. Comparison of the theoretical composition of a mock community obtained by mixing equal amounts of 27 plasmid DNAs containing a cytochrome *c* oxidase I insert from 27 strongyle species to determine the composition using deep amplicon sequencing. The frequencies obtained for individual species in theory and as measured are shown (A). All frequencies are compared with the expected frequency of 3.7% (B), which also shows the mean ± SD. Dotted lines are drawn at frequencies of 0% and the expected frequency of 3.7%. Genera are abbreviated as: *Cor., Coronocyclus; Cra.*, *Craterostomum; Cya., Cyathostomum; Cyc., Cylicocyclus; Cyd., Cylicodontophorus; Cys., Cylicostephanus; Gya., Gyalocephalus; Pet., Petrovinema; Pot., Poteriostomum; Str., Strongylus; Tri., Triodontophorus*.

### General description of deep amplicon sequencing data from field samples

The number of raw reads per sample obtained for equine-derived larval samples was in the range 676 – 331,284 (median 20,261) (S2 Table). Table 2 also contains metadata for all samples. After quality filtering, denoising, merging of forward and backward reads, and removal of chimeras, the number of reads was in the range of 396 – 19,3496 (median 13,733). Out of 124 samples, 70 samples had >10,000 reads, 24 had 5,000-10,000 reads, 21 had 1,000-5,000 reads, and nine had < 1,000 reads.

**Table 1.** Mean prevalence of individual strongyle species. Mean prevalence of individual strongyle species in samples from regularly (RT) and never treated (NT) equines

| Species | Regularly treated (N = 71) |  |  |  | Never treated (N = 50) |  |  |  | Comparison (reference = never treated) |  |  |  |  |
| --- | --- | --- | --- | --- | --- | --- | --- | --- | --- | --- | --- | --- | --- |
|  | n | Prevalence [%] | 95% CI |  | n | Prevalence [%] | 95% CI |  | Odds ratio | 95% CI |  | P<sub>unc.</sub> | P<sub>BH</sub> |
| *Coronocylus coronatus* | 37 | 52.1 | 40.7 | 63.3 | 26 | 52.0 | 38.5 | 65.2 | 1.00 | 0.48 | 2.09 | 0.990 | 0.990 |
| *Coronocylus labiatus* | 40 | 56.3 | 44.8 | 67.3 | 18 | 36.0 | 24.1 | 49.9 | 2.27 | 1.08 | 4.87 | 0.030 | 0.074 |
| *Coronocylus labratus* | 20 | 28.2 | 19.0 | 39.5 | 21 | 42.0 | 29.4 | 55.8 | 0.54 | 0.25 | 1.17 | 0.121 | 0.228 |
| *Coronocylus sagittatus* | 11 | 15.5 | 8.9 | 25.7 | 11 | 22.0 | 12.8 | 35.2 | 0.65 | 0.25 | 1.68 | 0.374 | 0.600 |
| *Craterostomum acuticaudatum* | 3 | 4.2 | 1.4 | 11.7 | 0 | 0.0 | 0.0 | 7.1 | n.a. |  |  | 0.198 | 0.352 |
| *Cyathostomum catinatum* | 59 | 83.1 | 72.7 | 90.1 | 32 | 64.0 | 50.1 | 75.9 | 2.73 | 1.17 | 6.56 | 0.020 | 0.058 |
| *Cyathostomum pateratum* | 34 | 47.9 | 36.7 | 59.3 | 6 | 12.0 | 5.6 | 23.8 | **6.52** | **2.60** | **19.08** | **<0.001** | **<0.001** |
| *Cyathostomum tetracanthum* | 6 | 8.5 | 3.9 | 17.2 | 0 | 0.0 | 0.0 | 7.1 | n.a. |  |  | 0.037 | 0.079 |
| *Cylicocyclus ashworthi* | 20 | 28.2 | 19.0 | 39.5 | 7 | 14.0 | 7.0 | 26.2 | 2.36 | 0.94 | 1.59 | 0.068 | 0.136 |
| *Cylicocyclus elongatus* | 1 | 1.4 | 0.2 | 7.6 | 3 | 6.0 | 2.1 | 16.2 | 0.25 | 0.01 | 2.19 | 0.218 | 0.3677 |
| *Cylicocyclus insigne* | 25 | 35.2 | 25.1 | 46.8 | 21 | 42.0 | 29.4 | 55.8 | 0.75 | 0.36 | 2.24 | 0.456 | 0.584 |
| *Cylicocyclus leptostomus* | 23 | 32.4 | 22.7 | 43.9 | 16 | 32.0 | 20.8 | 45.8 | 1.02 | 0.47 | 24.02 | 0.967 | 0.990 |
| *Cylicocyclus nassatus* | 63 | 88.7 | 79.3 | 94.2 | 23 | 46.0 | 33.0 | 59.6 | **8.94** | **3.67** | **0.39** | **<0.001** | **<0.001** |
| *Cylicocyclus radiatus* | 1 | 1.4 | 0.2 | 7.6 | 0 | 0.0 | 0.0 | 7.1 | n.a. | 0.07 | 6.90 | 0.587 | 0.722 |
| *Cylicodontophorus bicoronatus* | 2 | 2.8 | 0.8 | 9.7 | 12 | 24.0 | 14.3 | 37.4 | **0.10** | **0.01** | **0.39** | **<0.001** | **<0.001** |
| *Cylicostephanus bidentatus* | 0 | 0.0 | 0.0 | 5.1 | 1 | 2.0 | 0.4 | 10.5 | 0 |  |  | 0.413 | 0.575 |
| *Cylicostephanus calicatus* OTUI | 26 | 36.6 | 26.4 | 48.2 | 9 | 18.0 | 9.8 | 30.8 | 0.90 | 0.43 | 1.90 | 0.027 | 0.072 |
| *Cylicostephanus calicatus* OTUII | 37 | 52.1 | 40.7 | 63.3 | 15 | 30.0 | 19.1 | 43.8 | 0.39 | 0.18 | 0.83 | 0.017 | 0.054 |
| *Cylicostephanus calicatus* OTUIII | 0 | 0.0 | 0.0 | 5.1 | 1 | 2.0 | 0.4 | 10.5 | 0 |  |  | 0.413 | 0.575 |
| *Cylicostephanus goldi* | 45 | 63.4 | 51.8 | 73.6 | 31 | 62.0 | 48.2 | 74.1 | 1.06 | 0.50 | 2.25 | 0.877 | 0.935 |
| *Cylicostephanus longibursatus* | 70 | 98.6 | 92.4 | 99.8 | 42 | 84.0 | 71.5 | 91.7 | **11.70** | **2.00** | **301.95** | **0.004** | **0.018** |
| *Cylicostephanus minutus* OTUI | 28 | 39.4 | 28.9 | 51.1 | 21 | 42.0 | 29.4 | 55.8 | 0.90 | 0.43 | 1.90 | 0.780 | 0.891 |
| *Cylicostephanus minutus* OTUII | 32 | 45.1 | 34.0 | 56.6 | 34 | 68.0 | 54.2 | 79.2 | **0.39** | **0.18** | **0.83** | **0.014** | **0.0498** |
| *Cylicostephanus minutus* OTUIII | 33 | 46.5 | 35.4 | 58.0 | 28 | 56.0 | 42.3 | 68.8 | 0.69 | 0.33 | 1.42 | 0.311 | 0.498 |
| *Gyalocephalus capitatus* | 1 | 1.4 | 0.2 | 7.6 | 1 | 2.0 | 0.4 | 10.5 | 0.70 | 0.02 | 27.83 | 0.826 | 0.911 |
| *Petrovinema poculatum* | 4 | 5.6 | 2.2 | 13.6 | 13 | 26.0 | 15.9 | 39.6 | **0.18** | **0.05** | **0.55** | **0.002** | **0.011** |
| *Poteriostomum imparidentatum* | 18 | 25.4 | 16.7 | 36.6 | 5 | 10.0 | 4.3 | 21.4 | 2.97 | 1.08 | 9.80 | 0.035 | 0.079 |
| *Strongylus edentatus* | 5 | 7.0 | 3.0 | 15.4 | 30 | 60.0 | 46.2 | 72.4 | **0.05** | **0.02** | **0.15** | **<0.001** | **<0.001** |
| *Strongylus vulgaris* | 23 | 32.4 | 22.7 | 43.9 | 44 | 88.0 | 76.2 | 94.4 | **0.07** | **0.02** | **0.17** | **<0.001** | **<0.001** |
| *Triodontophorus brevicauda* | 2 | 2.8 | 0.8 | 9.7 | 3 | 6.0 | 2.1 | 16.2 | 0.47 | 0.05 | 3.18 | 0.431 | 0.575 |
| *Triodontophorus serratus* | 1 | 1.4 | 0.2 | 7.6 | 7 | 14.0 | 7.0 | 26.2 | **0.10** | **0.00** | **0.60** | **0.009** | **0.036** |
| *Triodontophorus tenuicollis* | 9 | 12.7 | 6.8 | 22.4 | 8 | 16.0 | 8.3 | 28.5 | 0.76 | 0.27 | 2.22 | 0.612 | 0.725 |

<sup>a</sup>95% confidence interval.
<sup>b</sup>Mid-p exact test. P<sub>unc.</sub> is uncorrected, P<sub>BH</sub> has been corrected for multiple testing using the method of Benjamini and Hochberg.

**Table 2.**
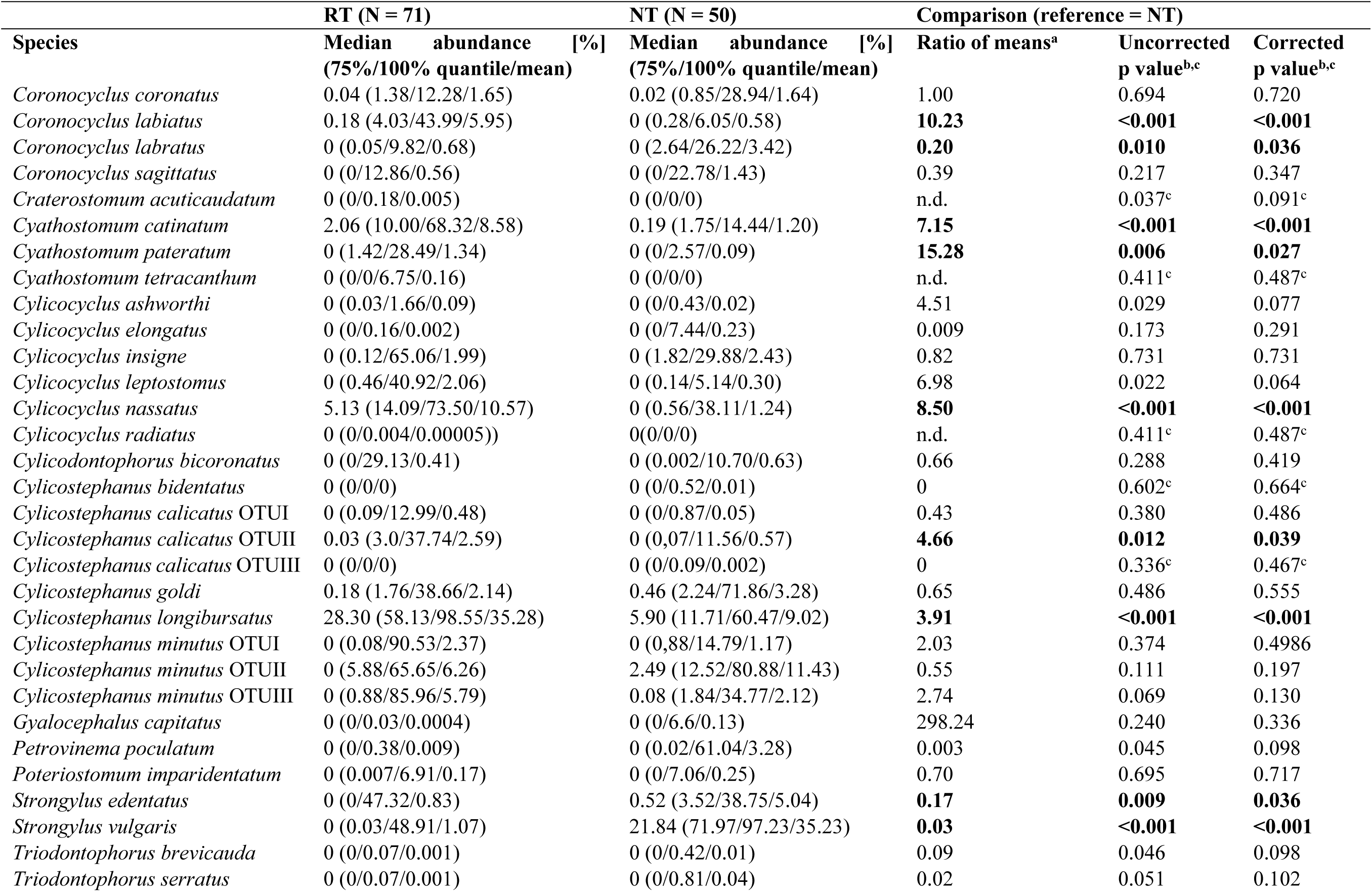
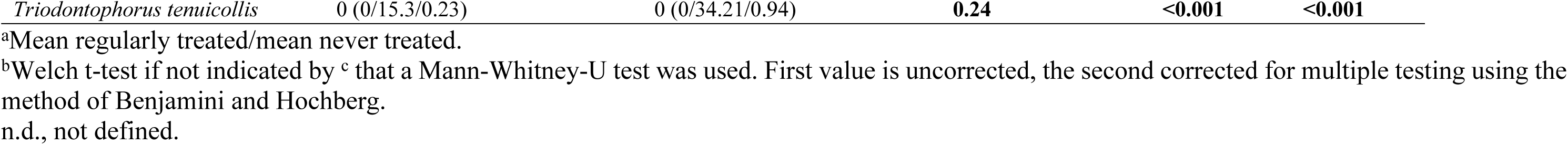
Relative median abundance of individual strongyle species. Relative median abundance of individual strongyle species in samples from regularly (RT) and never treated (NT) equines

### General description of species data obtained from field samples

In total, all 32 species, including three cryptic species in each of *Cys. calicatus* and *Cys. minutus*, which were included in the previously-developed sequence database (19) were detected in the field samples. The total number of filtered reads included was 3,030,149, representing 6,991 unique ASVs. Of these ASVs, 6,422 could be assigned to a species in the taxonomic database (19) using vsearch, while 565 ASVs (8.1%) remained unclassified. Unclassified ASVs that were not assigned to a particular species were present in 100 of 121 samples. The unclassified ASVs were represented by 394,838 reads (13.0% of the total read number). For individual samples, the percentage of unclassified reads ranged from 0 to 89.3% (median 1.2%, 75% quartile 9.4%). Although the two samples with the highest frequency of unclassified reads were from horses (a RT horse and a NT Przewalski’s horse from Ukraine), samples with a high frequency of unclassified reads were relatively rare in horse samples (including Przewalski’s horses), but occurred significantly more frequently in samples from domestic donkeys and kulans (Fig 3).

**Fig 3.**
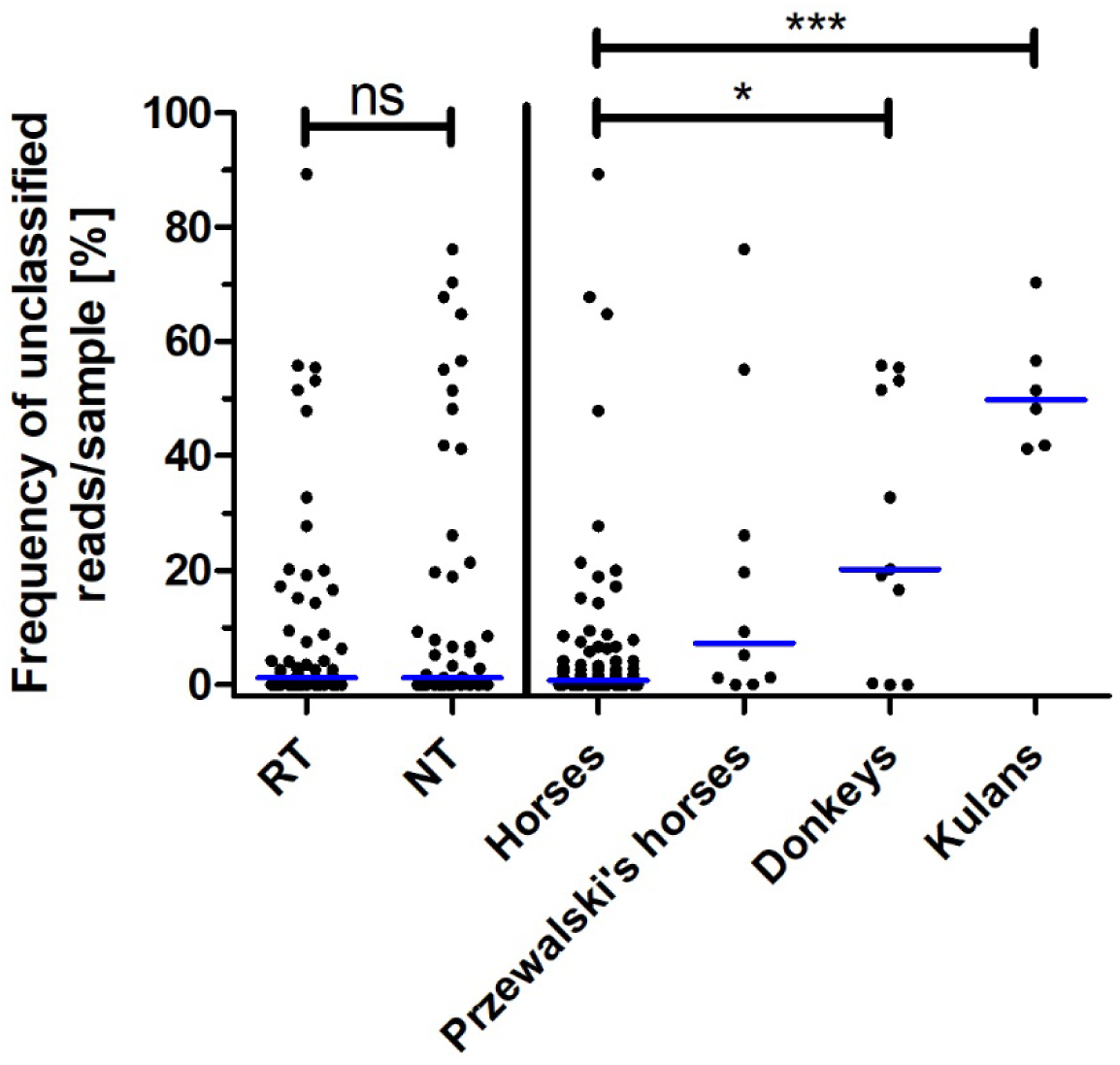
Frequency of unclassified reads that could not be assigned to a strongyle species using vsearch. Individual samples and medians are shown for samples from regularly (RT) or never treated (NT) equines and from domestic horses, Przewalski’s horses, donkeys, and kulans. For the latter, data from regularly and never-treated herds were pooled. Groups were compared using a Welch t test or Kruskal-Wallis test followed by Dunn’s post hoc test to compare all groups. Only significant comparisons are shown. *, p < 0.05; ***, p < 0.001.

Using rarefaction, it was evaluated whether sequencing depth was sufficient to estimate the species diversity of the samples. As shown in S1A Fig, most samples were close to saturation of ASVs detected, including samples for which read numbers were low (S1B Fig). There was a strong correlation (p < 0.0001; Pearson’s R = 0.67, Spearman ρ = 0.80) between the number of reads per sample and the number of detected ASVs (S2 Fig). According to linear regression, for each additional 1,000 reads, 2.9 additional ASVs were detected.

### Detection of individual species in field samples

Manual BLASTn analyses using COI from major non-strongylid parasitic nematodes of horses, i.e. *Tri. axei*, *Par. univalens*, *Str. westeri*, *Habronema* spp. and *Oxy. equi* and all ASV sequences detected in the dataset as subject revealed that none of the ASVs corresponded to these nematode species.

In total, all 32 species represented in the sequence database were identified in the field dataset. The strongyle species that were detected are listed in Table 1. This table compares the prevalence of these species between RT and NT animals using the NT animals as a reference group. After correcting for multiple testing, significant differences were observed for nine out of 31 species, with four species showing a higher prevalence in RT animals and five species showing a lower prevalence in RT animals. The large strongyle species *Str. edentatus* and *Str. vulgaris* showed the strongest difference with odds ratios of 0.05 and 0.07 (Table 1), respectively. This corresponds to 20.0- and 14.3-fold higher odds, respectively, to find an equine positive for *Str. vulgaris* or *Str. edentatus* in the NT group than for an equine in the RT group. Since *Str. equinus* was not included in the database, the complete set of ASVs detected in the equine samples were used as subject in a BLASTn analysis with the complete *Str. equinus* mitochondrial genome from GenBank (accession number NC_026868.1) as query. The best hits were ASVs assigned to *Str. vulgaris* and *Str. edentatus* with no evidence that any ASVs represented *Str. equinus*. Other species that occurred with significantly lower prevalence in RT equines were *Cyd. bicoronatus*, *Pet. poculatum*, and *Tri. serratus* (Table 1). In contrast, the odds of finding *Cys. longibursatus* increased 11.7-fold in samples from RT equines. Other strongyle species that were found significantly more often in animals from RT herds were *Cya. pateratum* and *Cyc. nassatus* (Table 1).

The relative abundance of all detected species is shown in Table 2, S3 Fig. and Fig 4 provide a comparison of the species detected in samples between RT and NT groups for all species and for those with significant differences in mean relative abundance, respectively. In RT horses, the most abundant species were *Cys. longibursatus*, *Cyc. nassatus*, *Cya. catinatum*, *Cys. minutus* OTUII, *Cys. minutus* OUTIII, and *Cor. labiatus.* For four of these species (*Cys. longibursatus*, *Cyc. nassatus*, *Cya. catinatum*, and *Cor. labiatus*), mean relative abundance was significantly higher in RT compared to the NT group (Table 2, Fig 5). In the NT group, the species with the highest mean relative abundance were *Str. vulgaris*, *Cys. minutus* OTUII, *Cys. longibursatus*, *Str. edentatus*, *Cor. labratus*, *Cys. goldi*, and *Pet. poculatum*. Three of these species (*Str. vulgaris*, *Str. edentatus*, *Pet. poculatum*) and *Tri. tenuicollis* were significantly more abundant in the NT in comparison to the RT group. Other species with significant differences in relative abundance between the RT and the NT groups were *Cya. pateratum,* and *Cys. calicatus* with higher abundance in the RT group.

**Fig 4.**
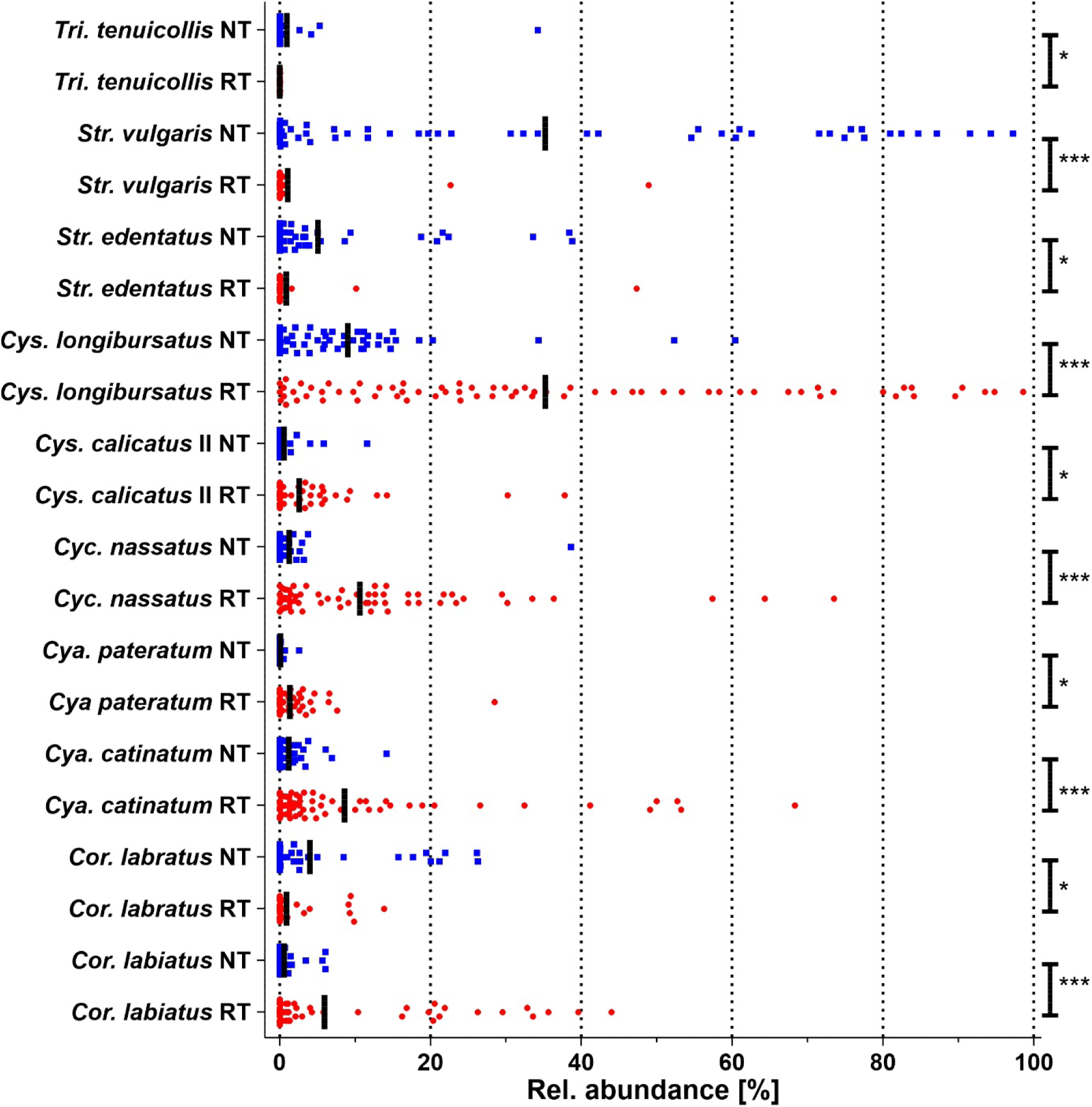
Relative abundance of identified strongyle species with significant differences between regularly and never treated equines. Treatment history is abbreviated as: RT (circles), regularly treated; NT (squares), never treated. Vertical lines represent medians. Data were compared between both groups using the Mann-Whitney U test, and p-values were corrected for multiple testing using the Benjamini-Hochberg approach. ***, p < 0.001; **, p < 0.01; *, p < 0.05. Genera are abbreviated as: *Cor., Coronocyclus; Cya., Cyathostomum; Cyc., Cylicocyclus; Cys., Cylicostephanus; Str., Strongylus; Tri., Triodontophorus*.

**Fig 5.**
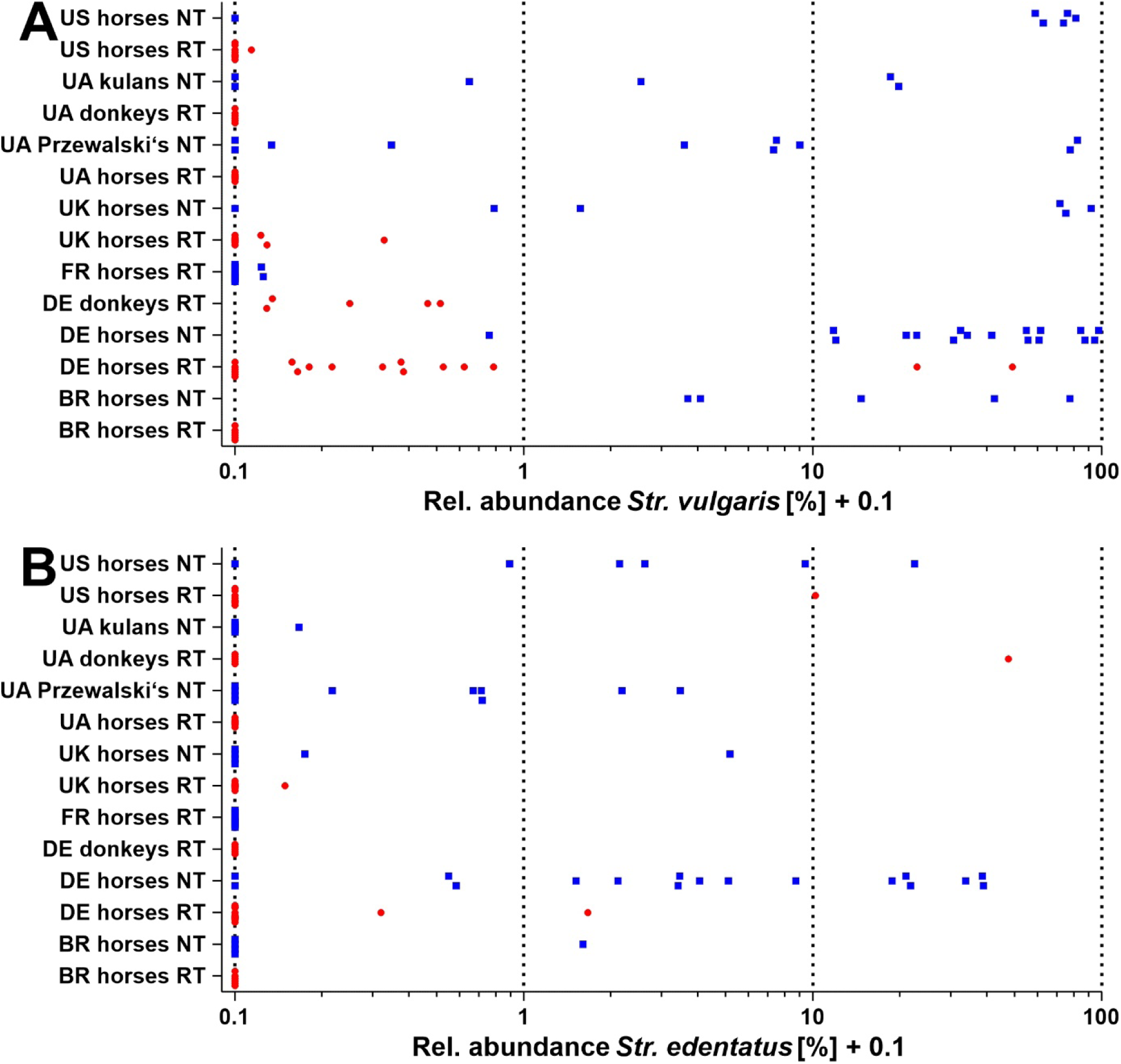
Relative abundance of *Strongylus vulgaris* (A) and *Strongylus edentatus* (B) in equine samples. Samples were collected in Brazil (BR), France (FR), Germany (DE), Ukraine (UA), the United Kingdom (UK), and the United States of America (US). Herds were either regularly treated (RT: circles) or never treated (NT: squares) in the last decades. Host species included domestic horses (horses), Przewalski’s horses (Przewalski’s), domestic donkeys (donkeys), and kulans. To allow usage of a logarithmic scale, 0.1% was added to all abundance values before plotting.

To help visualize the significant differences between RT and NT groups, data for parasites with significant differences are shown in Fig 4. Although some species occurred only in one of the two groups or with an extreme bias, the difference between both groups was often not significant because of very low relative abundance and occurrence of the species in a small number of equines. For those strongyle species for which differences in relative abundance were significant, the NT group was characterized by significantly higher mean relative abundance of the species *Str. vulgaris* (33.3-fold higher) and *Str. edentatus* (5.8-fold higher), *Cor. labratus* (5.0-fold higher), and *Tri. tenuicollis* (4.2-fold higher). In contrast, the RT group showed in significantly higher mean relative abundance of *Cya. pateratum* (15.3-fold higher), *Cor. labiatus* (10.2-fold higher), *Cyc. nassatus* (8.5 fold higher), *Cys. calicatus* OTUII (4.7-fold higher), and *Cys. longibursatus* (3.9-fold higher) (Table 2).

To determine if there was a correlation between the presence of *Strongylus* spp. and the five major cyathostomin species detected at significantly higher levels, the total relative abundance of *Strongylus* spp. and the five cyathostomin species *Cya. catinatum*, *Cya. pateratum*, *Cyc. nassatus*, *Cor. labiatus,* and *Cys. longibursatus* was calculated for each horse. The total relative abundances were then correlated, pooling data from RT and NT equines into a single dataset. S4 Fig. shows that there was a strong and highly significant negative correlation between the relative abundances of *Strongylus* spp. and those of Cyathostominae dominating in samples from RT equines.

To investigate whether the unexpectedly high prevalence of *Strongylus* spp. in RT equines was caused by a single or just a few herds of animals, the abundance of *Str. vulgaris* and *Str. edentatus* was plotted separately for each group (unique country, equine species, and treatment history) (Fig 5). To facilitate identification of samples with very low relative abundance of these parasite species, a logarithmic scale was chosen for the abundance. Very low relative abundance of *Str. vulgaris* below 1% occurred in 10/18 RT German horses (plus two with relative abundance >20%), 6/6 RT German donkeys, but also 2/10 RT French horses, 3/9 RT UK horses, and 1/9 RT USA horses. In contrast, *Str. vulgaris* was absent from RT horses and donkeys from Brazil and Ukraine. Low level of infections with *Str. edentatus* (<1% relative abundance) were detected in 1/18 RT German horses, 1/10 UK horses, 0/6 RT Ukrainian donkeys, and 1/9 RT USA horses. There were only three infections with *Str. edentatus* with a relative abundance >1% in the RT equines, and the parasite was completely absent from RT horses from Brazil, France, and Ukraine, and from RT donkeys from Germany. In contrast, relative abundances of *Str. vulgaris* and *Str. edentatus* in NT equines were much higher and frequently exceeded 1% and even 10% (Fig 5).

### Comparisons of α-diversity between RT and NT groups based on species and amplicon sequence variants

Since a considerable number of ASVs were not identified to species level, the α-diversity was analyzed based on both species and ASVs. Unclassified ASVs were considered to represent one additional species present in the samples. For all samples, the median number of species per individual equine was 11 (range 2 to 25). At the species level, species richness (observed richness and Chao1 index) and species diversity (Shannon and inverse Simpson index) showed no significant differences between RT and NT equines (Fig 6A). Unexpectedly, at the ASV level all four indices for richness (Fig 6B) and diversity (Fig 6C) showed significantly higher means for the RT compared to the NT group. Since RT is expected to decrease species richness and diversity, Chao1 and Shannon indices were also plotted for each group (same country, equine species, and treatment history) (S5 Fig.). For both indices, particularly high values were observed for RT equines from France and Germany. In the UK and the USA, species richness and diversity were also higher in RT than NT horses, while in Brazil and Ukraine, the opposite was observed (S5 Fig.). Since none of these groups was considered to be representative of the country or the host species, no formal statistical tests were conducted. Overall, the differences in species richness between RT and NT equines on the ASV level were apparently driven by the very high richness and diversity in samples from RT equines in Germany and France, but there was also an indication that there was a trend for the opposite relationship if these groups were excluded.

**Fig 6.**
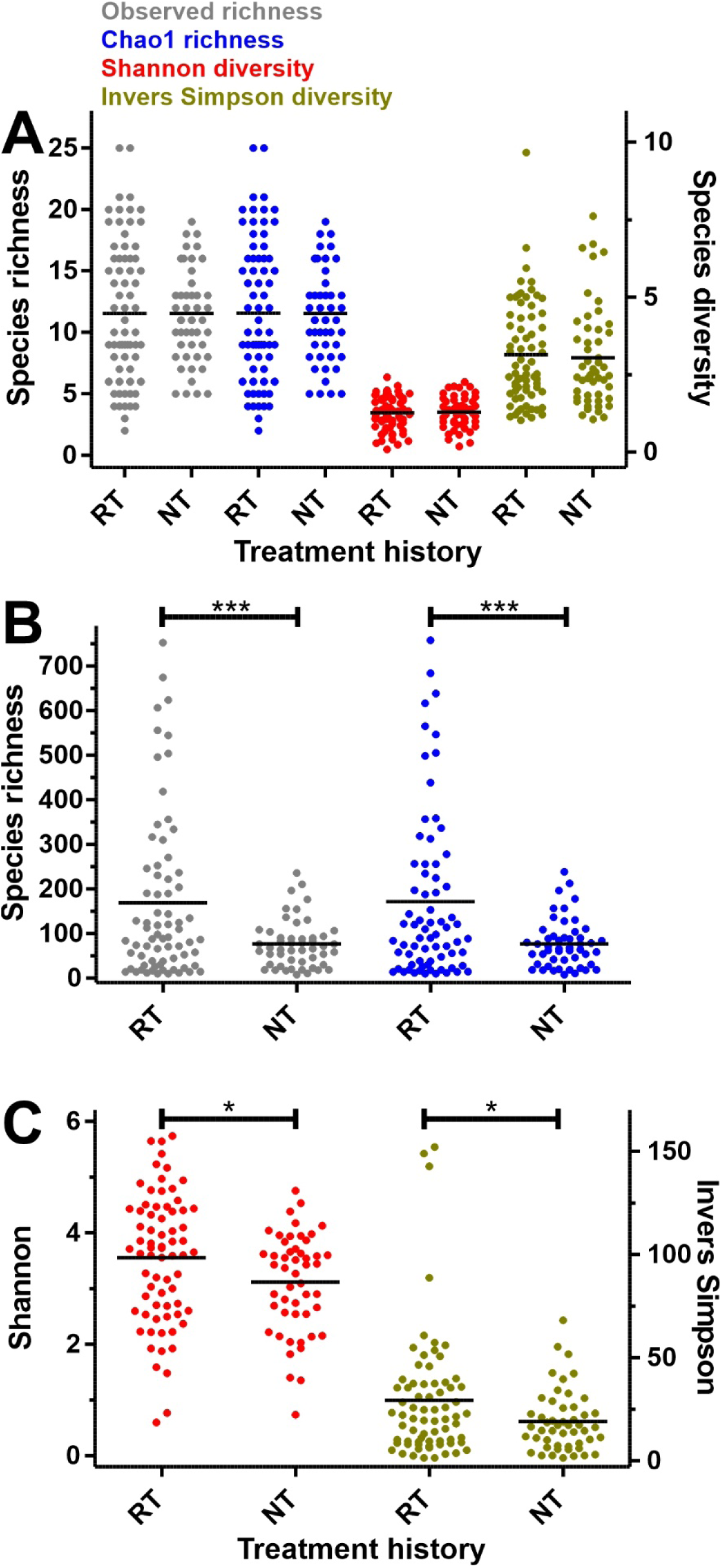
Species richness and diversity in regulary and never treated equine samples. Species richness and diversity at species (A) and amplicon sequence variant (B, C) levels in regularly (RT) and never treated (NT) equine samples. Species richness (observed richness and Chao 1 index) (A, B) and species diversity (Shannon and inverse Simpson index) (A, C) were calculated from species (A) and ASV data. Results for individual equine samples and means (horizontal lines) are shown. Data were compared between RT and NT groups using a Welch t test. ***, P < 0.001; *, p < 0.05.

### β-diversity of strongyle communities

From the frequencies of species in the individual equines, a Bray-Curtis and a Jaccard (based on binary presence/absence data) dissimilarity matrix were calculated based on abundance data. A PERMANOVA test (adonis2 of vegan package) (version 2.6-8) was used to detect if variables ‘treatment history’ (RT, NT), ‘country’, and ‘host species’ had an effect on dissimilarity between samples. As detailed in Tables S3 and S4 , all three variables had a highly significant effect on dissimilarity between samples. Since the data were considered to be not representative for the countries (only two study sites for several countries) and equine species (only one or two groups for kulans, Przewalski’s horses and donkeys), these variables were not further considered for analyses of β-diversity, which therefore focused only on the treatment history variable.

β-diversity was visualized using ordination based on non-metric multidimensional scaling (NMDS). The data show a clear separation between samples from equines derived from RT versus NT herds (Fig 7), with some overlap between the two groups. The NMDS plots for site (individual host sample) and species (parasite species) were plotted separately, since an overlay would not be readable due to the high number of data points. The NMDS plots show that species with significant differences in prevalence, abundance, or significant differences in the SIMPER analysis are clearly separated between RT and NT samples. Data based on the presence/absence of species in individual equine samples and a Jaccard dissimilarity matrix result in a very similar outcome with significant effects of the treatment history, the country of origin, and the equine species on β-diversity (S6 Fig.). In the NMDS plots, overlapping areas covered by samples from RT and NT horses were observed.

**Fig 7.**
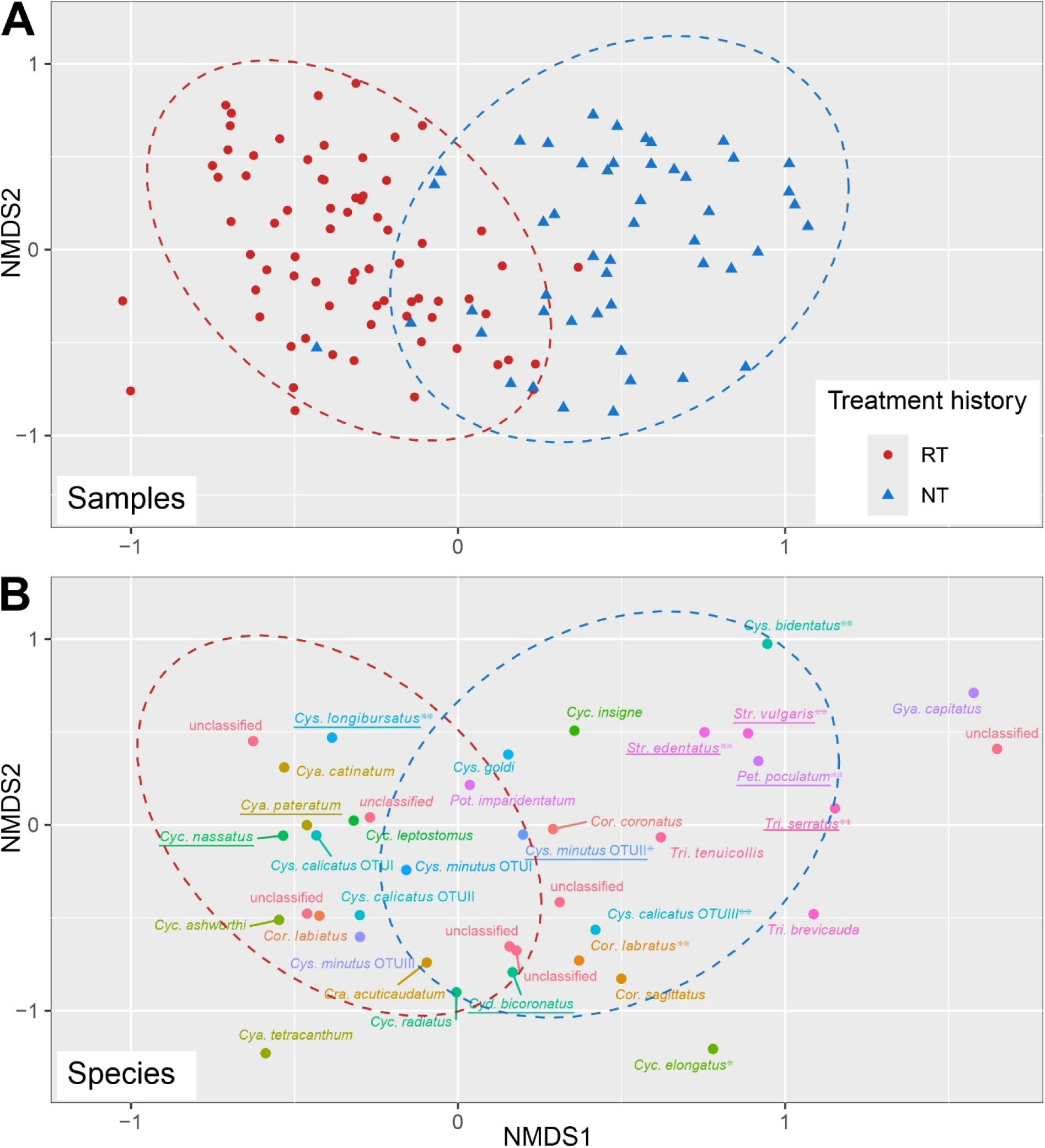
NMDS plot of strongyle species abundance in equine samples using a Bray-Curtis dissimilarity matrix. Non-metric multidimensional scaling (NMDS) plot based on the abundance of strongyle species identified in samples from individual equines using a Bray-Curtis dissimilarity matrix. Three-dimensional plots were generated since two-dimensional plots have stress values above 0.2. However, only dimensions 1 and 2 are shown as plots here. The plot for equine individual samples (study sites) is shown in (A), while the plot for parasite species is shown in (B) since an overlay of both plots was not readable. Ellipses in (A) show the areas in which 95% of samples from both treatment history groups are assumed to be located applying a t-distribution for the dissimilarity values. The ellipses for treatment history groups from (A) were transferred into the plot for parasite species in (B). This was only done to improve the comparability of plots in (A) and (B), which are on the same scale. Species with significant differences in prevalence and relative abundance are underlined or printed in bold, respectively. Significant differences between RT and NT groups according to the SIMPER analysis are indicated by asterisks. ***, P < 0.001; **, P < 0.01; *, P < 0.05. Genera are abbreviated as: *Cor., Coronocyclus; Cra.*, *Craterostomum; Cya., Cyathostomum; Cyc., Cylicocyclus; Cyd., Cylicodontophorus; Cys., Cylicostephanus; Gya., Gyalocephalus; Pet., Petrovinema; Pot., Poteriostomum; Str., Strongylus; Tri., Triodontophorus.* Multiple “unclassified” categories were included since samples were assigned to categories such as unclassified *Cylicocyclus*, unclassified *Strongylus*, etc.

A SIMPER analysis was conducted based on the Bray-Curtis dissimilarity matrix at species level to assess which strongyle species contributed most to the dissimilarity observed between RT and NT groups. Table 3 shows that the strongest contributions were made by *Str. vulgaris*, contributing 20.3% to the overall dissimilarity (more abundant in NT samples), and *Cys. longibursatus* and *Cys. minutus* OTUII, contributing 18.0% and 8.3% to the dissimilarity (more abundant in RT samples). Other species with a significant and considerable influence on dissimilarity were *Str. edentatus*, *Cor. labratus,* and *Pet. poculatum* (each of these species was more abundant in the NT group).

**Table 3.**
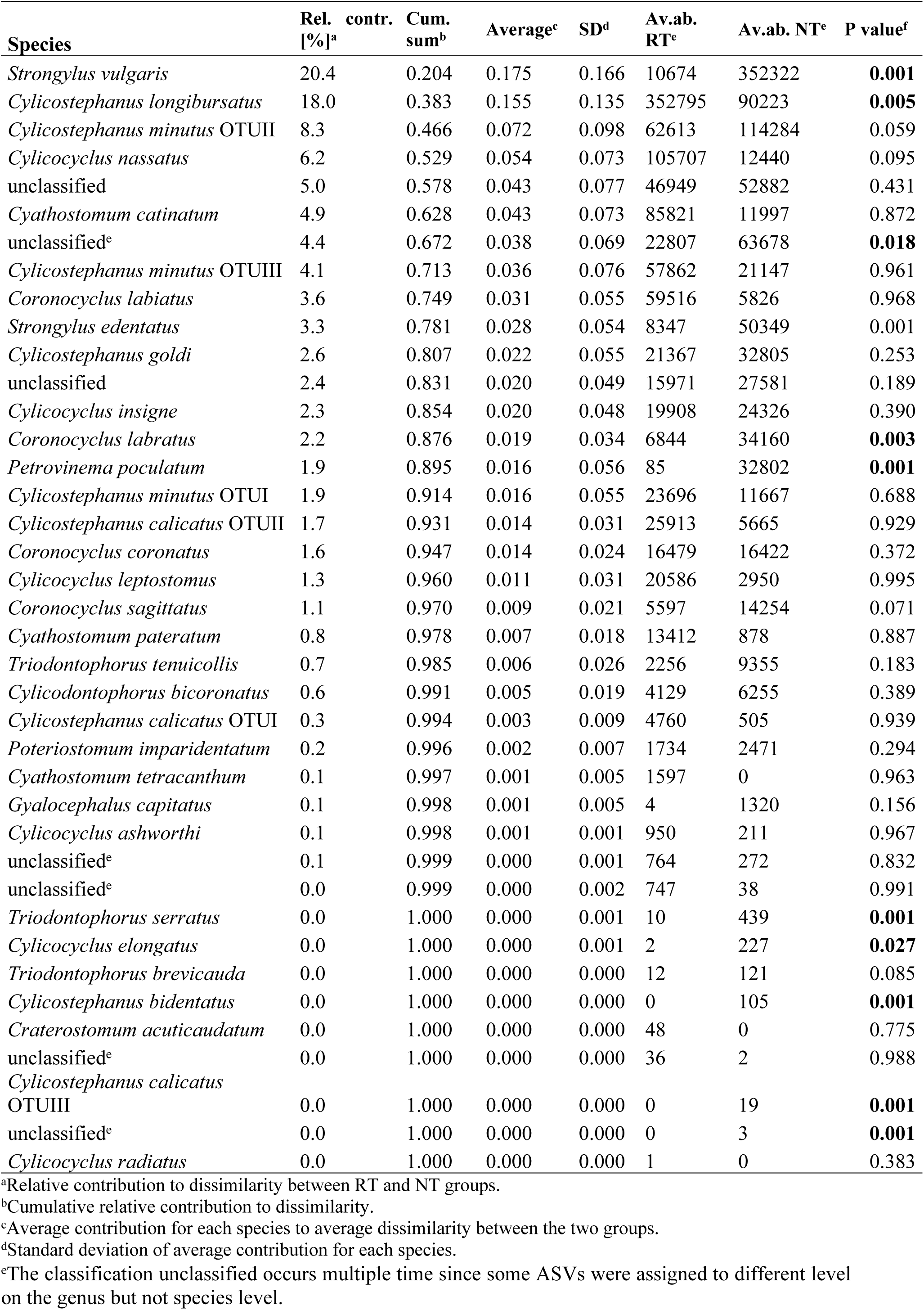
Dissimilarity between regularly treated and never treated equine groups. Contribution of strongyle species to dissimilarity between regularly treated (RT) and never treated (NT) groups based on SIMPER analysis data

| Species | Rel. contr. [%]<sup>a</sup> | Cum. sum<sup>b</sup> | Average<sup>c</sup> | SD<sup>d</sup> | Av.ab. RT<sup>e</sup> | Av.ab. NT<sup>e</sup> | P value<sup>f</sup> |
| --- | --- | --- | --- | --- | --- | --- | --- |
| *Strongylus vulgaris* | 20.4 | 0.204 | 0.175 | 0.166 | 10674 | 352322 | **0.001** |
| *Cylicostephanus longibursatus* | 18.0 | 0.383 | 0.155 | 0.135 | 352795 | 90223 | **0.005** |
| *Cylicostephanus minutus* OTUII | 8.3 | 0.466 | 0.072 | 0.098 | 62613 | 114284 | 0.059 |
| *Cylicocyclus nassatus* | 6.2 | 0.529 | 0.054 | 0.073 | 105707 | 12440 | 0.095 |
| unclassified | 5.0 | 0.578 | 0.043 | 0.077 | 46949 | 52882 | 0.431 |
| *Cyathostomum catinatum* | 4.9 | 0.628 | 0.043 | 0.073 | 85821 | 11997 | 0.872 |
| unclassified<sup>e</sup> | 4.4 | 0.672 | 0.038 | 0.069 | 22807 | 63678 | **0.018** |
| *Cylicostephanus minutus* OTUIII | 4.1 | 0.713 | 0.036 | 0.076 | 57862 | 21147 | 0.961 |
| *Coronocyclus labiatus* | 3.6 | 0.749 | 0.031 | 0.055 | 59516 | 5826 | 0.968 |
| *Strongylus edentatus* | 3.3 | 0.781 | 0.028 | 0.054 | 8347 | 50349 | 0.001 |
| *Cylicostephanus goldi* | 2.6 | 0.807 | 0.022 | 0.055 | 21367 | 32805 | 0.253 |
| unclassified | 2.4 | 0.831 | 0.020 | 0.049 | 15971 | 27581 | 0.189 |
| *Cylicocyclus insigne* | 2.3 | 0.854 | 0.020 | 0.048 | 19908 | 24326 | 0.390 |
| *Coronocyclus labratus* | 2.2 | 0.876 | 0.019 | 0.034 | 6844 | 34160 | **0.003** |
| *Petrovinema poculatum* | 1.9 | 0.895 | 0.016 | 0.056 | 85 | 32802 | **0.001** |
| *Cylicostephanus minutus* OTUI | 1.9 | 0.914 | 0.016 | 0.055 | 23696 | 11667 | 0.688 |
| *Cylicostephanus calicatus* OTUII | 1.7 | 0.931 | 0.014 | 0.031 | 25913 | 5665 | 0.929 |
| *Coronocyclus coronatus* | 1.6 | 0.947 | 0.014 | 0.024 | 16479 | 16422 | 0.372 |
| *Cylicocyclus leptostomus* | 1.3 | 0.960 | 0.011 | 0.031 | 20586 | 2950 | 0.995 |
| *Coronocyclus sagittatus* | 1.1 | 0.970 | 0.009 | 0.021 | 5597 | 14254 | 0.071 |
| *Cyathostomum pateratum* | 0.8 | 0.978 | 0.007 | 0.018 | 13412 | 878 | 0.887 |
| *Triodontophorus tenuicollis* | 0.7 | 0.985 | 0.006 | 0.026 | 2256 | 9355 | 0.183 |
| *Cylicodontophorus bicoronatus* | 0.6 | 0.991 | 0.005 | 0.019 | 4129 | 6255 | 0.389 |
| *Cylicostephanus calicatus* OTUI | 0.3 | 0.994 | 0.003 | 0.009 | 4760 | 505 | 0.939 |
| *Poteriostomum imparidentatum* | 0.2 | 0.996 | 0.002 | 0.007 | 1734 | 2471 | 0.294 |
| *Cyathostomum tetracanthum* | 0.1 | 0.997 | 0.001 | 0.005 | 1597 | 0 | 0.963 |
| *Gyalocephalus capitatus* | 0.1 | 0.998 | 0.001 | 0.005 | 4 | 1320 | 0.156 |
| *Cylicocyclus ashworthi* | 0.1 | 0.998 | 0.001 | 0.001 | 950 | 211 | 0.967 |
| unclassified<sup>e</sup> | 0.1 | 0.999 | 0.000 | 0.001 | 764 | 272 | 0.832 |
| unclassified<sup>e</sup> | 0.0 | 0.999 | 0.000 | 0.002 | 747 | 38 | 0.991 |
| *Triodontophorus serratus* | 0.0 | 1.000 | 0.000 | 0.001 | 10 | 439 | **0.001** |
| *Cylicocyclus elongatus* | 0.0 | 1.000 | 0.000 | 0.001 | 2 | 227 | **0.027** |
| *Triodontophorus brevicauda* | 0.0 | 1.000 | 0.000 | 0.000 | 12 | 121 | 0.085 |
| *Cylicostephanus bidentatus* | 0.0 | 1.000 | 0.000 | 0.000 | 0 | 105 | **0.001** |
| *Craterostomum acuticaudatum* | 0.0 | 1.000 | 0.000 | 0.000 | 48 | 0 | 0.775 |
| unclassified<sup>e</sup> | 0.0 | 1.000 | 0.000 | 0.000 | 36 | 2 | 0.988 |
| *Cylicostephanus calicatus* OTUIII | 0.0 | 1.000 | 0.000 | 0.000 | 0 | 19 | **0.001** |
| unclassified<sup>e</sup> | 0.0 | 1.000 | 0.000 | 0.000 | 0 | 3 | **0.001** |
| *Cylicocyclus radiatus* | 0.0 | 1.000 | 0.000 | 0.000 | 1 | 0 | 0.383 |

<sup>a</sup>Relative contribution to dissimilarity between RT and NT groups.
<sup>b</sup>Cumulative relative contribution to dissimilarity.
<sup>c</sup>Average contribution for each species to average dissimilarity between the two groups.
<sup>d</sup>Standard deviation of average contribution for each species.
eThe classification unclassified occurs multiple time since some ASVs were assigned to different level on the genus but not species level.

## Discussion

The data presented here demonstrate how deep amplicon sequencing is of value to the investigation of the effects of repeated anthelmintic treatments, which have potential to shape the composition of parasitic nematode communities in a multi-species complex. It was expected that the long-term application of anthelmintics would decrease the species diversity of strongyle communities due to repeated bottlenecking; this was recently described for two Ukrainian farms in a longitudinal study based on morphological identification of adult worms (45). In the present study, regular treatment of horses did not appear to lead to a reduction in species richness, and a lower degree of species diversity, as measured by assessment of ASVs, with significantly higher diversity and richness overall in the RT group compared to the NT group and no effect between both groups at species level. Repeated anthelmintic treatments reduced the number of *Strongylus* spp. detected in the present study and also in the morphological identification-based study in Ukraine (45). This could be due to the long phase (6-12 months) of larval development during the extraintestinal tissue migration of *Strongylus* spp. (46). Under field conditions, horses kept on conventional farms are likely treated with macrocyclic lactones active against migratory stages during the prepatent periods of these parasitic nematodes. Even a single annual treatment will considerably reduce the reproduction capacity of these species; this theory is supported by the reduced prevalence and clinical importance of large strongyles since macrocyclic lactones were introduced in the 1980s and 1990s (47–50).

In the present study, low levels of *Strongylus* spp. and a few other strongyle species were accompanied by a significantly higher abundance of several Cyathostominae species. *Cyc. nassatus*, *Cya. catinatum*, and *Cys. longibursatus*, three species identified as being the most prevalent and abundant worldwide (13) showed a strong bias towards samples from RT herds. All three species were among the top-six identified species in the list contributing to dissimilarity between RT and NT groups, but only for *Cys. longibursatus*, this contribution was significant. Although two other cyathostomin species showed a more significant bias towards RT samples, *Cya. pateratum* and *Cor. labiatus*; these species had an overall medium abundance in the dataset. That all three species, identified in a meta-analysis spanning five continents as highly abundant and prevalent, were also significantly more abundant in the RT group compared to the NT group here, indicates that regular treatment might have shaped the strongyle communities of domestic equines in recent decades worldwide (51, 52).

Interpretation of these results must acknowledge that the data on α- and β-diversity represent relative abundance of data on L3, with no information about the absolute number of adult and host larval stages available. The disappearance of some species, such as *Strongylus* spp., from the community will automatically lead to an increased relative abundance of the remaining species. This does not necessarily mean that the actual number of worms of these species has also increased. However, similar changes were observed for prevalence data, that β-diversity based on presence/absence data and that only the relative abundance of some, but not all, cyathostomin species was higher in the RT group suggest that anthelmintics have more complex effects than just strongly reducing the transmission of a few species with body tissue migration and long prepatent periods.

Noteworthy, the deep amplicon sequencing data unexpectedly showed that despite regular treatments, in some cases, *Str. vulgaris* and *Str. edentatus* remained present in equine strongyle communities at high prevalence but low relative abundance. If this is a general phenomenon, this is of major relevance for control strategies such as the ‘targeted-selective-treatment’ (TST) approach directed towards a reduced selection for anthelmintic resistance. If on a farm employing TST, the occurrence of *Strongylus* spp. is not recognized, e.g., due to diagnostic tools with low sensitivity, selective treatment strategies on farms with low abundance of *Strongylus* spp. have a risk that the abundance of these highly pathogenic parasites may increase in the absence of treatments. Findings of recent studies where significantly higher *S. vulgaris* infection risks were observed on farms claiming to use a selective treatment approach, as compared to strategic treatments, were published for Sweden, Denmark, and Germany (53–55). However, Halvarsson, Grandi (30) observed that the composition of strongylid communities in Swedish horses has remained relatively stable over several decades, despite the change in treatment regimen from strategic to TST practices since 1997. The study compared nemabiome data with data collected from morphologically identified adult worms from horses that had not been regularly dewormed and were not dewormed for at least a year before sampling (56). Since these horses cannot be considered as never treated, the strongylid composition may have been altered after the initial treatment. Consequently, the strongyle communities might have been shaped by general treatment practices rather than by selective or strategic treatment. In the present study, the large strongyle prevalence was particularly high in RT horses and donkeys from Germany, but infections with low relative abundance also occurred in several other countries. Larger cross-sectional studies are required to determine if this observed high prevalence of *Strongylus* spp. in RT horses was an exception or is representative. In the same geographic region from which the RT German horses originated, higher farm prevalence of *Str. edentatus* than *Str. vulgaris* has recently been reported using real-time PCRs (55). In the same project, a 21% prevalence of antibodies against a *Str. vulgaris* parasitic larvae (L4) antigen was reported at individual horse level. The prevalence of *Str. vulgaris* in the present study was 67%. Obviously, only 18 horses from two stables are not representative of the region. Nevertheless, the deep amplicon sequencing data presented here showed that a low relative abundance of *Strongylus* spp. in RT equines can occur, and this might lead to a high prevalence of antibodies against these parasites. The very low relative abundance, below 0.1-1% , observed in this study, might be associated with underdiagnosis of *Strongylus* spp. in relatively insensitive studies such as necropsy, but in particular after morphological discrimination of 100 L3 from coprocultures. The latter method has been shown to show only poor agreement with PCR analysis of the same samples (57). Comparison of real-time PCR and deep-amplicon sequencing methods for the detection of *Strongylus* spp. should be conducted to evaluate the validity of real-time PCR, which is much more likely to be conducted at single animal or farm level for routine diagnostic purposes than deep amplicon sequencing.

The nemabiome method used here, based on COI amplicons, has pros and cons compared to the more frequently used approach based on ITS-2 data. An obvious disadvantage of the COI-based method is the fact that PCR products are larger than the paired read length of the Illumina MiSeq. Due to the missing overlap of forward and reverse reads, it was not possible to correct raw sequences based on data from the reverse strand. This leads to higher error rates and higher numbers of ASVs caused just by experimental noise. Another disadvantage is that the COI primer pair used here is degenerated, while the NC1/NC2 ITS-2 primer pair is not. This might lead to different PCR efficacies for templates from different species; this was not observed for the species selected in the initial experiment comparing PCR efficacies. Advantages of the COI-based approach include the fact that the overlap in sequence identities between inter- and intra-species comparisons is small for COI but wide for ITS-2 sequences, with two frequently-occurring species having identical ITS-2 sequences (6, 19). Moreover, the alignment of COI sequences is reliable not only within the family Strongylidae but also in the whole order Strongylida and the phylum Nematoda, since alignments can be conducted based on relatively conserved protein sequences that exhibit rare insertions/deletions. In contrast, for ITS-2 sequences, it is often impossible to determine the best position of insertions/deletions in the alignment. This inaccuracy causes inaccuracy in species assignment. This also enables the identification of sequences not assigned to a taxonomic group to examine if they represent a reliable target sequence from a species in the taxon of interest (Strongylidae), and the Blast filtering step included in the bioinformatic pipeline uses this advantage to exclude ASVs that are not derived from the target taxon. This approach was successful since none of the sample sequences from other equine parasitic nematodes were identified, although the COI primer pair has been used before to amplify COI sequences from a clade III *Dracunculus* spp. nematode (58) and is expected to be useful across the phylum Nematoda. PCR efficacies are expected to have an effect on the results. The COI primer pair demonstrated comparable and high PCR efficacies for templates derived from different strongyle species. Moreover, the length of the PCR product was similar for COI, while ITS-2 PCR amplicons considerably differ in size, which might influence sequencing efficacy as Illumina shows bias towards short sequences (59).

Sequencing mock-communities is a standard control to demonstrate that a method can recover a defined community composition in terms of presence/absence and relative abundance. For equine strongyles, this is only possible to a limited extent because there are no monospecific isolates. To set up mock communities by mixing eggs or larvae from different species, multiple isolates from different species would be required; this is currently not possible. Mixing genomic DNA from species-identified adult worms as described by Courtot, Boisseau (44) is an alternative, but the quality of the DNA may differ between individuals and is difficult to quantify. Here, plasmid DNA with marker gene sequence inserts representing different species were used. The results of the mock community sequencing showed that most species in the mixture were detected by sequencing. The relative abundance of the species varied considerably. However, one species, *Cyc. elongatus*, was neither detected in the mock community nor in field samples, although it was previously amplified with the COI primer pair without Illumina adapters (19). Whether the species was simply not present because in recent decades *Cyc. elongatus* has been rarely detected in equine strongylid communities (45, 51, 52), or there were technical issues remains unknown.

There is a need to expand the equine nematode sequence database and add sequences from additional species. As detailed previously (19), most species with high or medium abundance identified in the recent meta-analysis (13) are included, and most gaps related to species of low or very low abundance. Herein, it was decided to avoid GenBank sequences to fill database gaps as multiple incongruencies were identified between the current dataset and those accessed in GenBank. One or the other sequence was assigned to the wrong species. Morphological differences between some of the species are small and extensive experience is required for reliable species identification, and thus, all specimens included in the current database were identified by Tetiana A. Kuzmina or Vitaliy A. Kharchenko, who together have decades of experience in the identification of strongylids by morphology (Lichtenfels et al., 2008). The approach used in the recent study by Diekmann et al. (2025b) was to identify, whenever possible, not only a single worm but multiple female and male specimens and obtain both ITS-2 and COI sequences from each specimen and resulted in high confidence that sequences in this database were assigned to the correct species. This is the reason why this database was employed as a reference here. Nevertheless, the database needs expansion since, for some species, only one or a few specimens were included, and other species are missing as no freshly collected and correctly preserved specimens were available. The database can detect most species occurring in domestic horses, but the number of reads that could not be assigned to a species was higher in domestic donkeys and kulans, suggesting that additional strongyle species, for which currently no sequence information is available, preferentially occur in these equids. It is well known that *Str. asini* and some Cyathostominae are specific parasites of donkeys and/or zebras, or at least only rarely occur in domestic horses (41). The large number of unclassified reads for donkeys and kulans suggests that there are more such species that occur only rarely in caballine but frequently in non-caballine equines, and that knowledge of the parasite species in the latter group is limited.

The data presented here confirmed the impact that anthelmintic treatment over decades has on the composition of strongyle communities in horses. Overall, regular anthelmintic treatment did not always lead to a decrease in species richness and diversity, which would have been expected if treatment led to frequent bottlenecking. However, since the most pathogenic species have the longest prepatent periods, regular anthelmintic treatment may lead to a replacement of pathogenic *Strongylus* spp. with other strongyle species. Important to note is the herein documented high prevalence but low relative abundance of *Str. vulgaris* and *Str. edentatus.* It has to be anticipated that in practice this will represent a challenge for the detection of large strongyle infections, which is particularly relevant regarding farms applying selective treatment strategies. The fact that identification of ASVs to species level was better for samples from horses than for donkey and kulan samples shows that the database, which currently contains domestic horse-derived samples, needs to be expanded using parasites specific for non-caballine equines.

## Materials and methods

### Selection of herds

In Brazil, all animals were of the Pantaneiro breed, which historically had limited human interference and became well-adapted to its environment, the Brazilian Pantanal. These NT horses (n=5) were raised extensively and on pasture and belonged to a large herd at Fazenda Campo Cyra, in the municipality of Rio Negro, in the state of Mato Grosso do Sul (MS).The samples were collected in July.2017 The Brazilian herd of RT horses (n=6) were sampled in August 2017 and came from the School Farm of the Federal University of Mato Grosso do Sul, in the municipality of Terenos (MS), where they were also raised on pasture but received regular anthelmintic treatments, especially with macrocyclic lactones.

In Nouzilly, France, samples from Welsh ponies were collected in May (n=8) and December (n=2) of 2019. The two horses sampled in December and the eight horses sampled in May had not been treated since 2017. Three horses were previously treated with pyrantel in October 2018.

In Germany, the NT horses (n=17) came from a feral horse (*Equus caballus*) group living largely under unmanaged conditions in the federal state North Rhine-Westphalia. This herd never received routine anthelmintic treatments. The samples were collected in July 2017. In contrast, a herd of RT horses (n=18) came from conventional horse farms originating from the federal states of Berlin and Brandenburg. The sample collection was in August 2017. The RT domestic donkey (*E. asini*) samples (n=5) were obtained from the federal states of Lower Saxony and Bavaria in Germany and were collected in April 2019. The treatment history and age of the individual animals was not recorded.

In Ukraine, the samples were collected in March 2016 from four species of equids kept in the “Askania Nova” Biosphere reserve – wild Przewalski’s horses (*E. ferus przewalskii*) (n=10) and Turkmenian kulans (*E. hemionus kulan*) (n=6) that were kept in semi-free conditions in large steppe enclosures and have never been treated with any anthelmintic and domestic horses (*Equus caballus*) (n=7) and donkeys (*E. asini*) (n=7) that were regularly treated with various anthelmintics twice a year during at least last two decades.

In Scotland, UK, samples were obtained in July 2017 from NT and RT horses. The NT UK samples (n=6) originated from a group of Konik ponies which were kept full-time on the Norfolk Broads, England, for conservation purposes. These horses were grazed extensively on Fen areas and had received limited treatments with anthelmintics. The regularly treated UK samples (n=10) originated from a yard on which the resident leisure horses of mixed breeds were previously treated with moxidectin in September 2016 (60).

In Kentucky, USA, samples from NT horses (n=7) were collected in May 2017 from a population of light-breed horses that had not been dewormed since 1979 (61). Samples from RT horses (n=7) were obtained from the research Shetland ponies’ herds “Population S” at the University of Kentucky (62). These horses were dewormed four times a year with ivermectin and ivermectin/praziquantel at the time of sample collection.

### Sample collection and larval cultures

All fecal samples were collected rectally or directly picked up from the ground after excretion to avoid contamination with soil nematodes. Larval cultures were set up from individuals. It was aimed to use more than 100 g fresh feces from each equine with at least 100 strongyle eggs per gram (epg) count. Loosened fecal material was incubated at 85% humidity and 25°C for 10 days without the addition of vermiculite or sawdust. The L3 were harvested from the fecal culture by filling the culture glasses with tap water, placing a petri dish on top of the glass followed by inverting the glass and filling the petri dish with tap water. Larvae that migrated from the fecal material into the petri dish were harvested, concentrated by sedimentation, purified using a Baermann funnel, and fixed in 70% ethanol. The larval numbers were not counted. The thus purified larvae were sent to the Institute for Parasitology and Tropical Veterinary Medicine in Berlin, Germany.

### DNA isolation and confirmation of presence of nematode DNA

Before DNA isolation, larvae were washed multiple times with PCR-grade water to remove the ethanol. DNA was isolated using the NucleoSpin Soil Mini kit (Macherey-Nagel, Düren, Germany) to remove potential PCR inhibitors.

### Library preparation

PCR’s were conducted with the primers COI_Nema_Fw and COI_Nema_Rv (63), which contain ILLUMINA adapters (4 nmole Ultramer® DNA Olígo, TruGrade®, Integrated DNA Technologies (IDT)) (S1 Table). PCR reactions contained 0.3 µM forward and reverse primers and 2 µl purified DNA in 25 µl 1× KAPA HiFi Hot Start ReadyMix (KAPABIOSYSTEMS, Cape Town, South Africa). The PCR program consisted of an initial denaturation at 95 °C for 180 s, followed by 35 cycles of denaturation at 98 °C for 20 s, annealing at 60 °C for 15 s, and elongation at 72 °C for 30 s, with a final extension at 72 °C for 60 s. PCR products from two identical reactions run in parallel were pooled, then purified on CleanNGS magnetic beads (CleanNA, Waddinxveen, The Netherlands) by mixing 20 µl PCR product with 32 µl bead suspension, followed by washing steps according to the manufacturer’s instructions and elution with 25 µl PCR-grade water. In a second PCR, indices were added to the PCR products using P7 and P5 primers with 8 bp indices obtained from biomers.net (Ulm, Germany) (S1 Table) . The PCR contained 0.5 µM of each primer and 10 µl purified PCR product from the first PCR round in 25 µl 1× KAPA HiFi Hot Start ReadyMix. The PCR started with denaturation at 98 °C for 45 s, followed by eight cycles of 98 °C for 15 s, 60 °C for 30 s, and 72 °C for 30 s. Finally, an extension was performed at 72 °C for 60 s. The second PCR product was purified on magnetic beads, combining 23 µl PCR product with 18.4 µl CleanNGS beads, and DNA was eluted in 20 µl PCR-grade water. DNA concentrations were determined using a Qubit 4™ fluorometer with dsDNA HS Assay Kit™ (Thermo Fisher Scientific, Waltham, MA, USA), and the quality of the libraries, in terms of size distribution, was evaluated using the DNA 5000 ScreenTape on a 4200 TapeStation (Agilent Technologies, Waldbronn, Germany).

To evaluate PCR efficacy for 12 strongyle species, plasmid DNA containing COI sequence inserts for different species was used as a template in the first PCR for the library preparation. The PCR reaction was modified by the addition of EvaGreen (Jena Bioscience) to a final concentration of 1 µM. The PCR was conducted in triplicate in a CFX96 real-time PCR cycler (Bio-Rad Laboratories, Feldkirchen, Germany), and fluorescence was recorded during the extension phase.

### Composition of mock communities

In the absence of reliable material to create mock communities from larval mixtures with defined composition, plasmid DNA containing COI sequences from 27 different operational taxonomic units (OTUs -including OTUs belonging to the same morphologically identified species) was mixed in equal amounts. DNA concentrations were measured using a Qubit 4™ fluorometer with dsDNA HS Assay Kit (Thermo Fisher Scientific, Waltham, MA, USA), then mixed to obtain a solution with 2 × 10^5^/µl plasmid copies. This mixture was used as a template for the same PCR’s that were used to construct libraries for field samples and then subjected to deep amplicon sequencing.

### Deep amplicon sequencing

Dual-indexed samples were pooled, test sequenced, and re-pooled according to read number for final sequencing. Sequencing was done with Illumina MiSeq using V3 600 sequencing kit (2x 300 cycles). Sequencing was performed at the Berlin Center for Genomics in Biodiversity Research.

### Bioinformatic analyses

Raw Next Generation Sequencing (NGS) reads were trimmed to remove primers using cutadapt version 4.4 (64) and untrimmed reads were discarded. Trimmed reads were then processed using dada2 version 1.26.0 (65). After quality control, a truncation length of 200 bp and 180 bp was selected for forward and reverse reads, respectively. For the truncated reads, dada performed error learning and denoising before the denoised forward and reverse reads were paired and chimeras removed using the removeBimeraDenovo function in consensus mode. As the amplicon length significantly exceeds the combined read lengths, pairing was performed in “justConcatenate” mode, where reads are not overlapped but concatenated with a 10 Ns separator string. The resulting sequences were aligned against the known variant database using blastn version 2.14 (66) and rejected unless both reads matched with at least 80% identity and 90% query coverage. Assignment to known variants was performed with strict thresholds of 100% identity and coverage within a 30 bp window relative to the 5’ and 3’ ends of the known variant. For species assignment, the sintax mode of vsearch version 2.27.0 (67) as used with a sintax_cutoff of 0.8. For downstream analyses, samples with less than 500 total sequence counts and sequences. Alpha and beta diversity measures and figures were generated using the R package phyloseq version 1.44 (68).

BLASTn was used to determine whether the dataset contained reads from non-strongyle parasitic nematodes of horses. For this purpose, the *Trichostrongylus axei* (GenBank accession no NC_013824.1), *Parascaris univalens* mitochondrial genome (NC_024884), *Strongyloides papillosus* mitochondrial genome (NC_028622), *Habronema muscae* partial COI (FJ471583) and *Oxyuris equi* mitochondrial genome (NC_027190) were used as the query. A detailed description of the bioinformatic pipeline implemented as a Snakemake (69) workflow and bash script, is available in a FigShare repository (https://doi.org/10.6084/m9.figshare.28840316.v1).

### Statistical analyses

PCR amplification efficacy was calculated from real-time PCR amplification plots for each replicate using LinRegPCR (version 2021.1) and default conditions (70–72). PCR efficacies were plotted for each species and a One-way ANOVA without post-hoc test was performed using GraphPad Prism 5.03 to evaluate if there were differences in efficacy between species. The relative abundance of individual species was compared between RT and NT groups using the Welch t test in GraphPad. Prevalences of individual species were compared between RT and NT groups using the mid-p exact test as implemented in the oddsratio.midp() and tab2by2.test() functions in the epitools package (version 0.5-10.1) in R 4.3.3. The p-values obtained for all detected species were corrected for multiple testing using the Benjamini-Hochberg method (5% false-discovery-rate) using the p.adjust() function in R. Pearson and Spearman correlations between variables were calculated in GraphPad Prism.

## Acknowledgements

The authors thank the horse owners for their valuable support and help. The authors also thank the research training group GRK 2046 (Graduierten-Kolleg 2046) for stimulating discussions that substantially improved the manuscript.

## Data availability statement

All data are available in the figures, tables within the manuscript and Supporting Information files. Sequencing data are available at NIH’s Sequence Read Archive (SRA) under BioProject accession number: PRJNA1248057. The bioinformatic workflow is available from the Figshare data repository: https://doi.org/10.6084/m9.figshare.28840316.v1.

Data are available under the terms of the <u>Creative Commons Attribution 4.0 Internationallicense</u> (CC-BY 4.0).

## A Supporting information

**S1 Fig. Evaluation of saturation of amplicon sequence variant (ASVs) numbers using rarefaction.** The number of species that were detected was plotted over the number of reads included in the subsample for all samples (A) and for samples for which only low numbers of reads were available (B).

**S2 Fig. Correlation of post-BLAST filter reads and ASVs per sample**. Correlation of the number of reads per sample after BLAST filtering and the number of ASVs detected in the same sample. The results of linear regression and Pearson’s correlation coefficient R are indicated.

**S3 Fig. Relative abundance of all detected strongyle species from regularly and never treated equines.** Treatment history is abbreviated as: RT (circles), regularly treated; NT (squares), never treated. For significant differences between species see Table 2. Genera are abbreviated as: *Cor., Coronocyclus; Cra.*, *Craterostomum; Cya., Cyathostomum; Cyc., Cylicocyclus; Cyd., Cylicodontophorus; Cys., Cylicostephanus; Gya., Gyalocephalus; Pet., Petrovinema; Pot., Poteriostomum; Str., Strongylus; Tri., Triodontophorus*.

**S4 Fig. Correlation between relative abundance of *Strongylus* spp. with relative abundance of the five Cyathostominae species.** Correlation between relative abundance of *Strongylus* spp. with relative abundance of the five Cyathostominae (Cyath.) species dominating in regularly treated (RT) equines (*Cyathostomum catinatum*, *Cyathostomum pateratum*, *Cylicocyclus nassatus*, *Coronocyclus labiatus*, and *Cylicostephanus longibursatus*). Individual samples were labeled according to the assignment to the groups of RT and never treated (NT) equines. The correlations and linear regression analysis were conducted on pooled data from both groups.

**S5 Fig. Species richness as Choa1 and diversity as Shannon index.** Species richness as Choa1 (A) and diversity as Shannon index (B) calculated from amplicon sequence variant data for strongyle species from different countries and equine species from herds with different treatment histories. Results for individual equine samples and means (horizontal lines) are shown. Samples were collected in Brazil (BR), France (FR), Germany (DE), Ukraine (UA), the United Kingdom (UK), and the United States of America (US). Herds were either regularly treated (RT) or never treated (NT) in the last decades.

**S6 Fig. NMDS plot of strongyle species abundance in equine samples using a Jaccard dissimilarity matrix.** Non-metric multidimensional scaling (NMDS) plot based on the abundance of strongyle species identified in samples from individual equines using a Jaccard dissimilarity matrix calculated from binary (presence/absence) data. Three-dimensional plots were generated since two-dimensional plots have stress values above 0.2. However, only dimensions 1 and 2 are shown as plots here. The plot for equine individuals (study sites) is shown in (A), while the plot for parasite species is shown in (B) since an overlay of both plots was not readable. Ellipses in (A) show the areas in which 95% of samples from both study history groups are assumed to be located, applying a t distribution for the dissimilarity values. The ellipses for study history groups from (A) were transferred into the plot for parasite species in (B). This was done to improve the comparability of plots in (A) and (B), which are on the same scale. Species with significant differences in prevalence and relative abundance are underlined or printed in bold, respectively. Significant differences between RT and NT groups according to the SIMPER analysis are indicated by asterisks. ***, P < 0.001; **, P < 0.01; *, P < 0.05. Genera are abbreviated as: *Cor., Coronocyclus; Cra.*, *Craterostomum; Cya., Cyathostomum; Cyc., Cylicocyclus; Cyd., Cylicodontophorus; Cys., Cylicostephanus; Gya., Gyalocephalus; Pet., Petrovinema; Pot., Poteriostomum; Str., Strongylus; Tri., Triodontophorus.* Multiple “unclassified” categories were included since samples were assigned to categories such as unclassified *Cylicocyclus*, unclassified *Strongylus*, etc.

**Table S1. Barcode primer & ILLUMINA adapter for the 1.PCR and index-primer for the 2.PCR step**

**Table S2. Metadata and number of raw reads per sample**

**Table S3. Bray-Curtis distance matrix.** Effects of equine treatment history, equine species (horses, Przewalski’s horses, donkeys, kulans) and country of origin on dissimilarity between samples using a PERMANOVA analyses with 9999 permutations based on a Bray-Curtis distance matrix on the parasite species level.

**Table S4. Jaccard distance matrix.** Effects of equine treatment history, equine species (horses, Przewalski’s horses, donkeys, kulans) and country of origin on dissimilarity between samples using a PERMANOVA analyses with 9999 permutations based on a Jaccard distance matrix on binary (presence/absence) data on the parasite species level.

